# Physiological adaptation to high irradiance in duckweeds depends on light habitat niche and is ecotype and species-specific

**DOI:** 10.1101/2023.06.27.546714

**Authors:** Kellie E. Smith, Laura Cowan, Beth Taylor, Lorna McAusland, Matthew Heatley, Erik H. Murchie

## Abstract

Duckweeds are free-floating aquatic organisms with species ranging from 2 mm-10 mm, where each plant is a single leaflike structure. Recognized as an emerging food crop, their fast growth rates offer potential for cultivation in closed systemsHowever the majority of available duckweed clones lack information regarding habitat origin and physiology. We describe a novel UK collection derived from low light (dLL) or high light (dHL) habitats and profiled for growth, photosynthesis and photoprotection (Non Photochemical Quenching, NPQ) responses. Multiple ecotypes of three *Lemna* species and one ecotype of *Spirodela polyrhiza*, were grown under low light (LL:100 μmol m^-2^ s-1) and high light (HL:350 μmol m^-2^ s-1). We found species and ecotypic variation in photosynthesis acclimation. Duckweeds grown under HL exhibited lower growth rate, biomass, chlorophyll and quantum yield of photosynthesis. In HL-compared to LL, carotenoid de-epoxidation state and NPQ were higher whilst photosystem II efficiency (ϕPSII) and chla:b ratios were unchanged. Interestingly dLL plants showed relatively stronger acclimation to HL compared to dHL plants: These ecotypes achieved faster growth in HL: by area and colony gain, higher carotenoid levels and less degradation of chlorophyll. We conclude that adaptation to local habitat among ecotypes strongly affects performance under controlled conditions.

## Introduction

Duckweeds are miniature, aquatic plants that can match the fastest growth rates of higher plants, with population doubling times < 24 hours and potential high dry weight production of 106 tonnes ^-1^ ha^-1^ year^-1^ (Cui & Cheng, 2015; Ziegler *et al*., 2015; Michael *et al*., 2020). Species are widely distributed across lentic ponds and swamps and flowing water bodies of streams and canals. Consisting of simplified units of tiny, reduced leaf-stem structures (fronds) detaching to form independent clones, asexual reproduction allows rapid colonization and full coverage of water surface habitats. The 36 duckweed species have great phenotypic diversity within five genera including larger and rooted *Spirodela, Landoltia, Lemna*, and the shrunken and rootless *Wolffia and Wolffiella*. Duckweeds have world-wide market potential as human or animal feeds: they grow fast in soil-less cultivation, have global distributions and offer a promising source of dietary protein in the form of essential amino acids comparable to soybean (Cheng & Stomp, 2009). Duckweeds are also a source of high starch, fibre, micro and macronutrients (Appenroth *et al*., 2017; Yahaya *et al*., 2022). Historically used in Asian cooking, there is also growing interest in duckweeds for vertical farming and as a live plant food for extended space exploration missions and to fill gaps in longer-term astronaut nutrition (Smith *et al*., 2009, Smith *et al*., 2015). (Appenroth *et al*., 2018; Stewart *et al*., 2020; Polutchko *et al*., 2022; Nguyen *et al*., 2023).

As miniature plants, duckweeds may have little control over growing position, maintaining a static position in the environment, often lodged at water edges or passively motile, carried with the water flow or by water-dwelling organisms through an array of light environments. Duckweeds are tolerant of extremely low light conditions from grates and drains where light does not penetrate, to vast duckweed carpets in expansive open lakes such as Titicaka in Peru and Bolivia, showing tolerance of growth in full light and high temperature (Vargas-Cuentas & Roman-Gonzalez, 2019). Although successful worldwide colonization strategies suggests natural variability of photosynthetic apparatus to challenging light climes, this has not been explored extensively in duckweeds (Ceschin *et al*., 2018; Paolacci *et al*., 2018a; Stewart *et al*., 2020).

Plants have a multitude of mechanisms that allow efficient adaptation to diverse intensities and wavelengths of light in natural habitats, caused by e.g., season, cloud cover and tree and shrub cover. Whilst light stimulates high photosynthesis under plant optimal conditions, high irradiance can cause long term damage to photosynthetic processes (photoinhibition) often in combination with other stresses e.g., low or high temperature. Plants can modify area and width of leaves, re-arrange composition of light harvesting complexes, photosystem stoichiometry, pigment composition, photosynthetic rates per unit leaf area and increase production of carotenoids and other counter-acting antioxidants (Anderson *et al*., 1995; Maxwell *et al*., 1995; Murchie & Horton, 1997, Foyer & Harbinson, 1999; García-Plazaola *et al*., 2004, Walters, 2005, Poorter *et al*., 2006). Photoacclimation processes typically support high photosynthetic capacity in high light and more efficient light harvesting and quantum yield under low light. This is commonly observed using light response curves for gas exchange and / or electron transport showing quantum yield and the saturation point of photosynthesis which can vary e.g. 300-500 µmol m^2^ s^-1^ for temperate annuals to around 1000 µmol m^2^ s^-1^ in rice growing in the tropics approximately (Murchie & Horton, 1997; Zhao *et al*., 2017). Acclimation to high light is also associated with photoprotective processes such as non-photochemical chlorophyll fluorescence (NPQ) which quenches excess excitation energy as heat and helps to prevent photoinhibition. NPQ consequentially downregulates quantum yield of photosynthesis and can restrict plant growth in prolonged dynamic light exposure in natural conditions (Kromdijk *et al*., 2016; Ruban, 2016, 2017; Pniewski & Piasecka-Jędrzejak, 2020).

Great species variation exists in nature in terms of the ability to acclimate to light conditions which can be linked to growth strategy (Murchie & Horton, 1997, 1998; Demmig-Adams *et al*., 2012; Burgess *et al*., 2023). Cultivar/accession variation in photosynthetic properties has been shown in traditional food crops and their close relatives including wheat and rice (e.g. McAusland *et al*., 2020; Cowling *et al*., 2022). It follows that both ecotype and species variation in duckweed selection should be considered. Variation in growth and changes in thylakoid pigments and photosynthesis in response to light has started to be considered in single *Lemna* clones of different species (Paolacci *et al*., 2018a; Stewart *et al*., 2020) whilst it seems *Lemna* species contain an enrichment in light harvesting proteins compared to *Spirodela* species (An et al., 2018). However, there has not been a comprehensive analysis of variation in acclimation of photosynthesis to light in duckweed species and ecotypes and it has not been linked with adaptation to habitat irradiance.

Duckweeds may present an interesting challenge for photoacclimation due to their floating habit across diverse sites, unusual frond anatomy which includes large air spaces for floating, high stomatal conductance, rapid colony formation and minimal photoassimilate export to vasculature. The growth rate is usually assessed by clonal colony expansion rates rather than progressive (3D) canopy development and tropic responses toward light seen in other macrophytes. In duckweeds, growth rate showed species and ecotypic variation in stable light conditions (Ziegler *et al*., 2015), however light adaptation should be placed in the context of growth rate to better understand how photosynthetic adaptations can help to achieve the high growth rates in these species.

Understanding the links between growth, photosynthesis, light tolerance and nutritive potential can also aid in the selection of duckweeds as candidates for new food applications and for optimized vertical farming trials and space missions. For instance, carotenoids involved in light adaptation are desirable in animal feed (Ekperusi *et al*., 2019) and human food as they have benefits to human health (Eggersdorfer & Wyss, 2018).

The objectives of this study are to determine natural variation in adaptation to light in duckweeds from wild sites with specific light parameters and quantify the environmental acclimation and tolerance of duckweeds to light spatially and temporally. We hypothesise that growth rate will vary between ecotypes, with photosynthetic and photoprotective properties linking to originating habitats. Those hailing from high light environments are expected to show stronger responses of photoacclimation to high light when grown under controlled environmental conditions.

## Materials and methods

### Collection of duckweed ecotypes and measurements of environmental parameters

24 duckweed isolates were collected in May 2020 from UK sites between latitudes of 49.9 and 53.9 ° and longitudes of −0.29 and −5.19 ° (see Table S1, Fig 1A). Between 10-20 individuals from each site were collected into sealed tubes of water. If mixed species were visible across the site, individuals were taken of each type into different sample tubes. These samples were stored at ambient temperature with natural daylengths until return to the laboratory, where species were confirmed using genotyping. Site KS06 was a special case where two *L. minuta* KS06A and KS06B cultivars were sourced however KS06B came from a drain excluding light, and was measured separately to test for potential extreme light adaptive differences.

**Figure 1.**
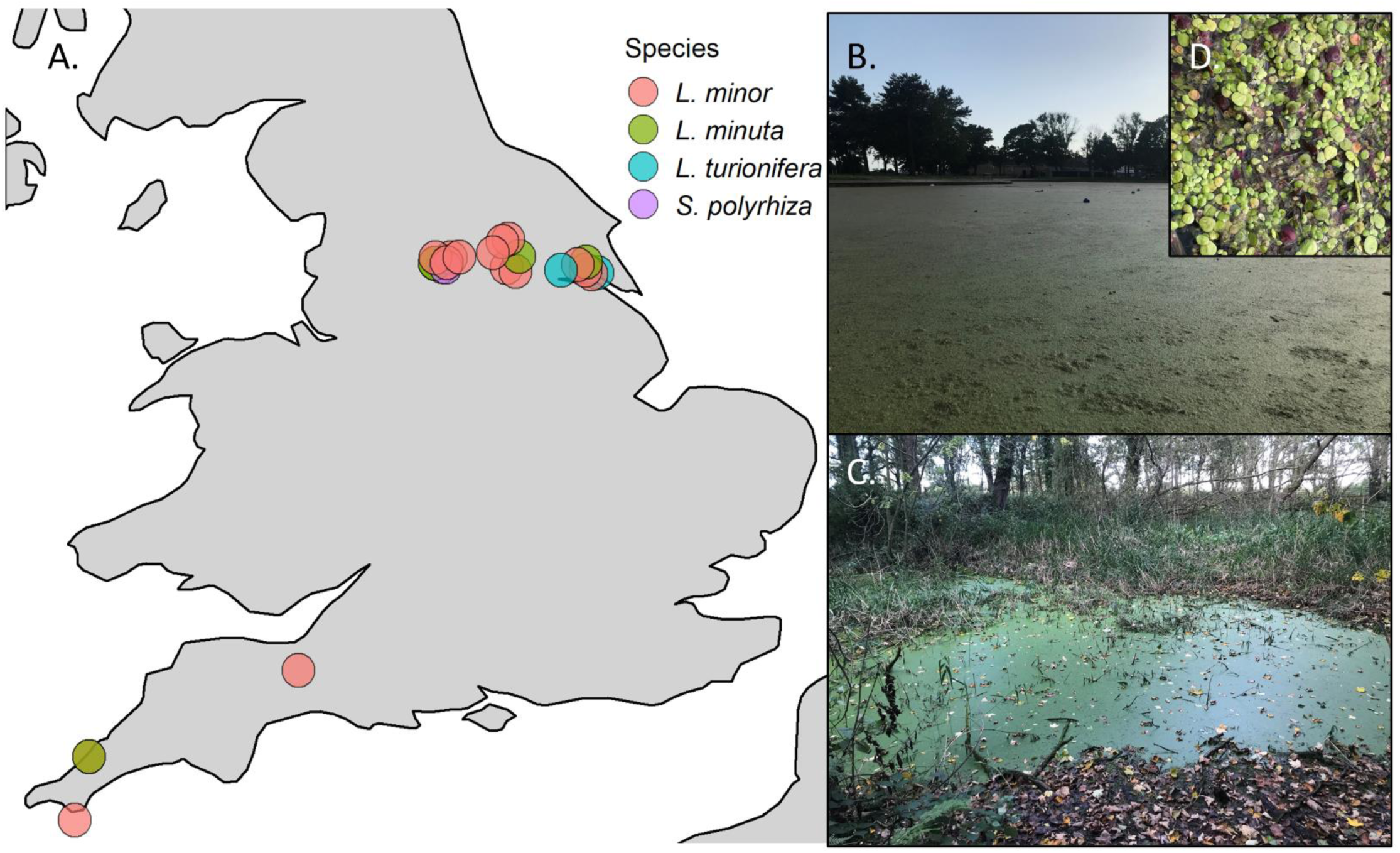
UK collection sites and high duckweed coverage found at high and low light sites. *A.* Map of collection sites of duckweeds in this study, plotted by longitude, latitude coordinates (S1. Table). Colored circles represent species groupings 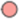 *L. minor*, 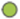 *L. minuta*, 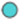 *L. turionifera*, 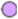 *S. polyrhiza* (as determined in Fig S5). *B.* High duckweed coverage in an open pond: KS12 in Bradford, England, high light (dHL) site. *C.* High duckweed coverage in a pond with steep banks and high tree cover: site KS25 in York, England representative of low light (dLL) site. Inset. D. Close up of purple coloration in *S. polyrhiza* at KS12 dHL site.

Nineteen duckweed sites were visited to collect environmental data over a two-day period in March 2021 and 2022 and July and October in 2020, 2021 and 2022 to monitor variation across the seasons of Spring, Summer and Autumn.

Duckweed coverage scores were estimated on site between 0 and 5 determined by surface coverage. Three photos from above were taken per site with a Canon 650D camera suspended on a camera boom. A white reference and scale was provided for each photo, level with the water surface. An umbrella was used to minimize light interference through water reflection. Images processed as follows: three representative areas of 5×4 cm rectangles were selected to determine duckweed coverage average and variability per site. Images were split into red, green and blue stacks and the blue stack used with the threshold scale to best match the original photos. Percentage coverage was quantified using Fiji open-source software (Schindelin *et al*., 2012). Coverage data and locational coordinate data for duckweed sites is provided in (Fig S1).

Light intensity (maximum PPFD, photosynthetic photo flux density intensity) was measured above 10 cm or up to 1 m (as close as possible to a) water source using a 400 – 700 nm light meter (LICOR, Li-250A) with an attached quantum sensor head. Using a handheld spectrometer (LICOR, LI-1500) total PFD, PPFD and the ratio of light wavelengths making up the PFD (380-780 nm) were split and recorded as PFD-UV (380-400 nm), PFD-B (400-500 nm), PFD-G (500-600nm), PDF-R (600-700 nm), and PFD-FR (700-780 nm) in µmol m^-2^ s^-1^ and dominant (λd) and peak (λp) wavelengths (in nm) recorded at each site. Dominant wavelength is defined as the color perceived and the peak wavelength as the highest intensity wavelength recorded per site. All measurements were taken three times over the 20-minute period of the visit and maximum recorded. All light intensity and spectrum variables (total = 45) were used together to group duckweed sites as dLL or dHL habitats using distance analysis and K-means clustering to split into two main groups by site similarity (Fig S2.). Distance matrices were computed using Manhattan, Euclidean and Mikowski and showed good consistency for two main groupings and site similarities. To measure variety in light environments, proportions of spectral quality at each timepoint was calculated as proportion of each individual spectral region/total light PFD x 100 per site to give percentages and R:FR was calculated as ratio of R to FR from raw values from each site.

Time, weather (cloud cover) and atmospheric and water temperatures were noted across sites to account for variability across the two-day periods. Climate data was collected for each longitude latitude combination using Bioclimatic variables extracted from worldclim.org using R package Bioclim (Serrano-Notivoli *et al*., 2022) and altitudes obtained relative to sea level using Google Earth Pro. The raw and grouped light data sets are summarised in (Fig S2.) and other environmental data shown in Fig S3.

### Maintenance of duckweed ecotypes

To sterilise, wild duckweeds were treated with 0.5% sodium hypochlorite in well plates (Greiner bio-one, Cellstar) for 1-2 mins. Time of treatment was dependent on size of duckweed and visible inward bleaching rates, leaving a green meristematic pocket and then dipping in Milli-Q water 18 MΩ to recover. Multiple individuals from a site were either designated A or B based on either size (different species) and cultured separately. Sterile colonies were grown in individual flasks containing N media. Duckweed stocks were grown in GEN-1000 cabinets (Conviron, Winnipeg, Canada) with light provision at 50 µmol m^-2^ s^-1^ PPFD using broad spectrum white LED lights providing 16:8 days with a ramp of light intensity at the start and end to represent sunset and sunrise. A temperature cycle of 22 / 18 °C day and night, and relative humidity maintained at 60%. N media was prepared with Milli-Q water 18 MΩ and consists of KH_2_PO_4_ (0.15 mM), Ca(NO_3_)_2_ (1 mM), KNO_3_ (8 mM), MgSO_4_ (1 mM), H_3_BO_3_ (5 µM), MnCl_2_ (13 µM), Na_2_MoO_4_ (0.4 µM), FeEDTA (25 µM) as described in Appenroth *et al*., 1996). N media was autoclaved before use at 121° C for 20 mins. Each week duckweeds were re-sterilised with 0.5% sodium hypochlorite with dipping into sterile Milli-Q water and placed into fresh flasks containing new media to build up sterile stock populations.

### Determining duckweed species

Each of the sterile 24 duckweed ecotype stocks were harvested into small populations containing 5-20 individuals and frozen in liquid nitrogen. Individuals were ground using a Tissuelyser II (Qiagen) and DNA extracted using DNAeasy Plant kit (Qiagen). DNA quantification was performed using dsDNA HS assay (ThermoFisher Scientific) and Qubit 2.0. Individual library preparations and short read sequencing using Illumina HiSeq 2500 platform sequencing was performed by Novogene, Cambridge, UK. Species identification was called using a genomic pipeline as described in (Monnahan *et al*., 2019) and available at (https://github.com/mattheatley/ngs_pipe). Datasets for *Lemna* and *Spirodela* ecotypes published in the sequence read archive (SRA) were also included as references for species groups. Briefly, all individuals were aligned to a common reference *L. minor* 7720 using GATK version 3.5 with several filtering steps to obtain SNP variation. The allele frequencies of ecotypes were examined by PCA using adegenet package (version 2.1.3) on R (Jombart, 2008) with independent classifications based on phenotypic observations also performed (i.e. size of duckweed and presence or absence of seed formation). Briefly, all individuals were aligned to a common reference *L. minor* 7720 using GATK version 3.5 with several filtering steps to obtain SNP variation.

### Controlled growth conditions for light acclimation experiments

GEN-2000SH cabinets (Conviron, Winnipeg, Canada) installed with broad spectrum white LED lights were used to provide growth light treatments. Six months after collection and cabinet domestication at 50 µmol m^2^ s^-1^ light, UK accessions were sub-cultured to around 15-20 individuals from each accession into individual flasks and continued to be grown long days 16/8 hr at 22/18 °C day night temperatures. Accessions were either placed for two weeks at low light (LL: 100 µmol m^2^ s^-1^) (individual ecotypes n = 24) or in 150 µmol m^2^ s^-1^ for one week to acclimate to the intermediate light condition and then transferred to high light (HL: 350 µmol m^2^ s^-1^) for a further week (n=24), to acclimate to conditions (Stewart *et al*., 2020). The experiment with each light program then ran constantly for up to six weeks with a one-hour light and temperature ramp up and down to simulate sunrise and sunset. N media was changed weekly to maintain sterile populations with high nutrient density. Temperature and relative humidity were recorded using Datalogger (TinyTag Ultra 2, Gemini data loggers) in addition to the cabinet sensors. Light intensity and spectra were measured using light sensor (LICOR, LI-1500) (Fig 2A).

**Figure 2:**
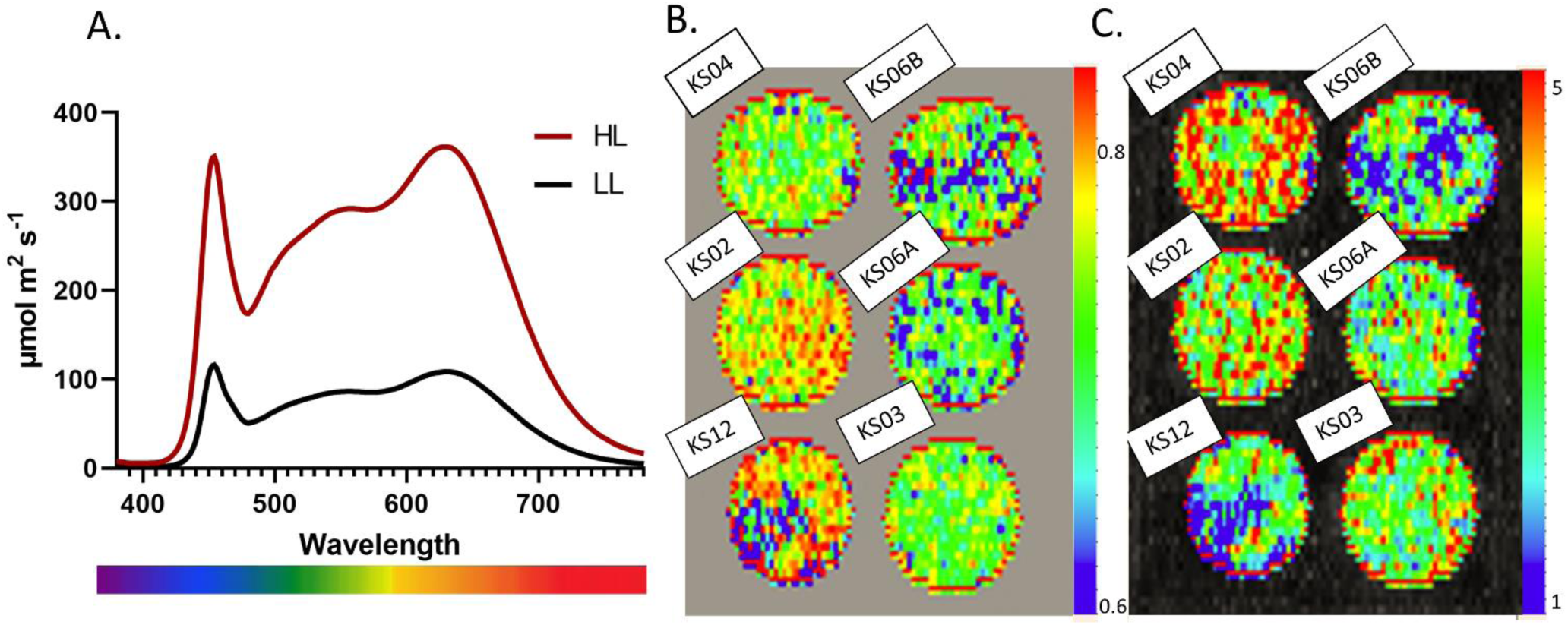
*A.* spectra for individual wavelengths in PPFD region provided in two duckweed light growing conditions. HL – PPFD 350 µmol m^2^ s^-1^ and LL – PPFD 100 µmol m^2^ s^-1^. False colour images applied to pixels corresponding to duckweed cultivar populations within well plates as measured by chlorophyll fluorescence (B). *F*_v_ */ F*_m_ and (C.) NPQ measured by a Fluorcam. Individual accessions are labelled per well and grown at HL. Scales represent ranges of measurement values 0.6-0.83 for *F*_v_ */ F*_m_ in the dark and values of 1-5 for NPQ at maximum light.

### Growth rate measurements

Single three-frond colonies from light level stock populations were added to individual conical flasks with 100 ml N media on day 0 (n=24) ecotypes for each HL and LL treatment. Growth rate was measured for each ecotype in each condition until ∼95% surface coverage was achieved, or for up to six weeks in slower growing ecotypes. N media was replaced each week to maintain high nutrient levels. Relative growth rate (RGR) was measured by colony gain, by counting the number of colonies in each flask, in each condition, every seven days. Col RGRlog and RGRlog by area gain were calculated from raw data (see Fig S5). RGRlog by area gain was derived from total green area (mm^2^) measured using a digital Nikon D5100 camera and an imaging pipeline for quantification (Ware *et al*., 2023). RGR difference by area and colony gain between treatments was obtained by mean growth in HL – mean growth in LL and proportion change by RGR difference/mean growth in LL x 100. Total fresh weight biomass (FW) per flask (normalized by weight of starter colony) was measured after 6 weeks (T42). Fresh biomass per flask was harvested and snap frozen in liquid nitrogen before being freeze dried and weighed to obtain FDW.

### Chlorophyll fluorescence

Populations of each ecotype were grown for four or six weeks in each light level. Populations of each ecotype-treatment combination were used to fill the surface of clear plastic six-well plates containing 3 ml N media. Chlorophyll fluorescence imaging was carried out according to (McAusland *et al*., 2019) and layout in six well plates as shown in Fig. 2B. Duckweed plates were imaged using a closed chlorophyll fluorescence imager (800C Fluorcam, Photon System Instruments, Brno, Czech Rep.) after a 1hr dark adaption. White LEDs with actinic light provided a saturating pulse set at 4500 µmol m^2^ s^-1^ for 0.8 secs to measure *F*_v_/*F*_m_. Then a rapid stepwise light response programme was used increasing from 0, 20, 130, 245, 365, 480, 600, 710, 830, 950, 1050 µmol m^2^ s^-1^ PPFD (referred to as L1 – L11 level steps corresponding to light levels) with a saturating pulse applied at the end of each step. Numeric averages were exported for each ecotype-treatment replicate using the in-built software (Fluorcam 7, Photon System Instruments, Brno, Czech Rep.). The following parameters were extracted from the protocol: maximum photosystem II (PSII) efficiency (*F*_v_/*F*_m_), PS II operating efficiency (*F*_q_’/*F*_m_’) or ϕPSII and NPQ (*F*_m_–*F*_m_′)/*F*_m_′) (non-photochemical quenching i.e. the photoprotective dissipation as heat loss) at each light level (Murchie & Lawson, 2013).

### Pigment extraction and analysis

For spectrophotometry, five mg freeze-dried duckweed tissue was ground in 80% 1.5 ml acetone using a TissueLyser II (Quigen) at 24 Hz/sec for 4 mins and cell debris pelleted. Extracted supernatant was further diluted by 3.5 ml 80% acetone to give a total volume of 5 ml. Absorbance was recorded for chl a at 663 nm, chl b at 646 nm, carotenoids at 470 nm and absorbance at 750 nm as a correction turbidity factor using a UV/Visible spectrophotometer (Ultrospec 2100 pro, Amersham biosciences). Total chlorophyll and carotenoids as mg/g duckweed were calculated following (Porra *et al*., 1989).

### Pigment extraction and analysis by HPLC

For carotenoid HPLC analysis, tissue was rapidly frozen in liquid nitrogen at mid day. 0.8 g was ground in 2 ml 100% acetone (HPLC grade) in low light and centrifuged at 10,000 rpm at 4°C for 2 mins. The supernatant was filtered through a 13 mm diameter 0.2 µm polytetrafluoroethylene (PTFE) syringe filter (Whatman GmbH, Dassel, Germany) into a 1.5 ml amber Eppendorf and stored at −80°C. Pigment separation was performed by reverse-phase HPLC as described in (Färber *et al*., 1997) using a Dionex BioLC HPLC system (Sunnyvale, California, US) with LiChrospher^®^ 100 RP-18 (5 µm) column (Merck, Darmstadt, Germany). Xanthophylls: violaxanthin, zeaxanthin, antheraxanthin and other carotenoids lutein, neoxanthin and β-carotene and chl a and chl b were detected using 447 nm wavelength, shown in the chromatogram (see Fig S10A). Individual carotenoids were expressed as % of the total of carotenoids and the de-epoxidation state (DEPs), total xanthophyll pool (XC) calculated as described in (Färber *et al*., 1997).

### Experimental design and statistical analysis methods

Experiments were repeated on five separate occasions giving five independent set of growth experiments. From each of these, three biological replicates were used for each ecotype-treatment combination for chlorophyll fluorescence measurements giving up to 15 replicates. Pigment extraction by spectrophotometry was performed for up to four replicates of ecotype-treatment combinations from each independent experiment, maximum n= 20. For HPLC analysis, 14 ecotypes grown in HL and LL were used, forming one rep of each ecotype-treatment combination. Significance of accession, species, treatment and derived environmental light (dHL or dLL) was performed on each growth rate parameter (RGRlog area, Col RGRlog, FW, FDW) using all observation data using Two-way ANOVA and Welch’s T-tests. The RGR difference between treatments were determined by HL – LL growth data for each experimental repeat. The differences HL – LL for averaged data for NPQ, **ϕ**PSII and *F*_v_/*F*_m_ and pigment contents (total chlorophyll a, b and carotenoids) and HPLC data were used to find differences between treatments. All data manipulation and analysis was performed in R (v3.6.3) using Rstudio (v1.2.5) with packages ggplot2 (Gómez-Rubio, 2017), corrplot from (McKenna *et al*., 2016), FactoMineR (Lê *et al*., 2008) and factoextra (Kassambara & Mundt, 2020).

## Results

Collection was made from sites with a range of light levels, from full scale open ponds and canals, to ditches and locations under bridges and trees (Fig 1B, 1C). Full coverage was visible in both light and shade locations and surface coverage varied between sites and seasons (S1. Table). Environmental light data (Fig. S2) was correlated with coverage measurements of 19 sites across the seasonal-year time points (Fig. S1). In Autumn, coverage was significantly negatively correlated with UV light, with an R2 of −0.54 *P value* = 0.02 (Fig S4). There were no other correlations between light and coverage.

Duckweed sites were successfully classified as derived high light (dHL) or low light (dLL) using distance analysis and K-means clustering to form two groups using all measured environmental light variables of light intensity PPFD and all spectral components of light measurements at sites across timepoints (Fig 3, and data summarised in Fig 4 with statistical test results in Table S2. See also ‘environmental collection’ in methods). Eleven collection sites were determined as low light sites and eight from high light sites (Fig 3).

**Figure 3:**
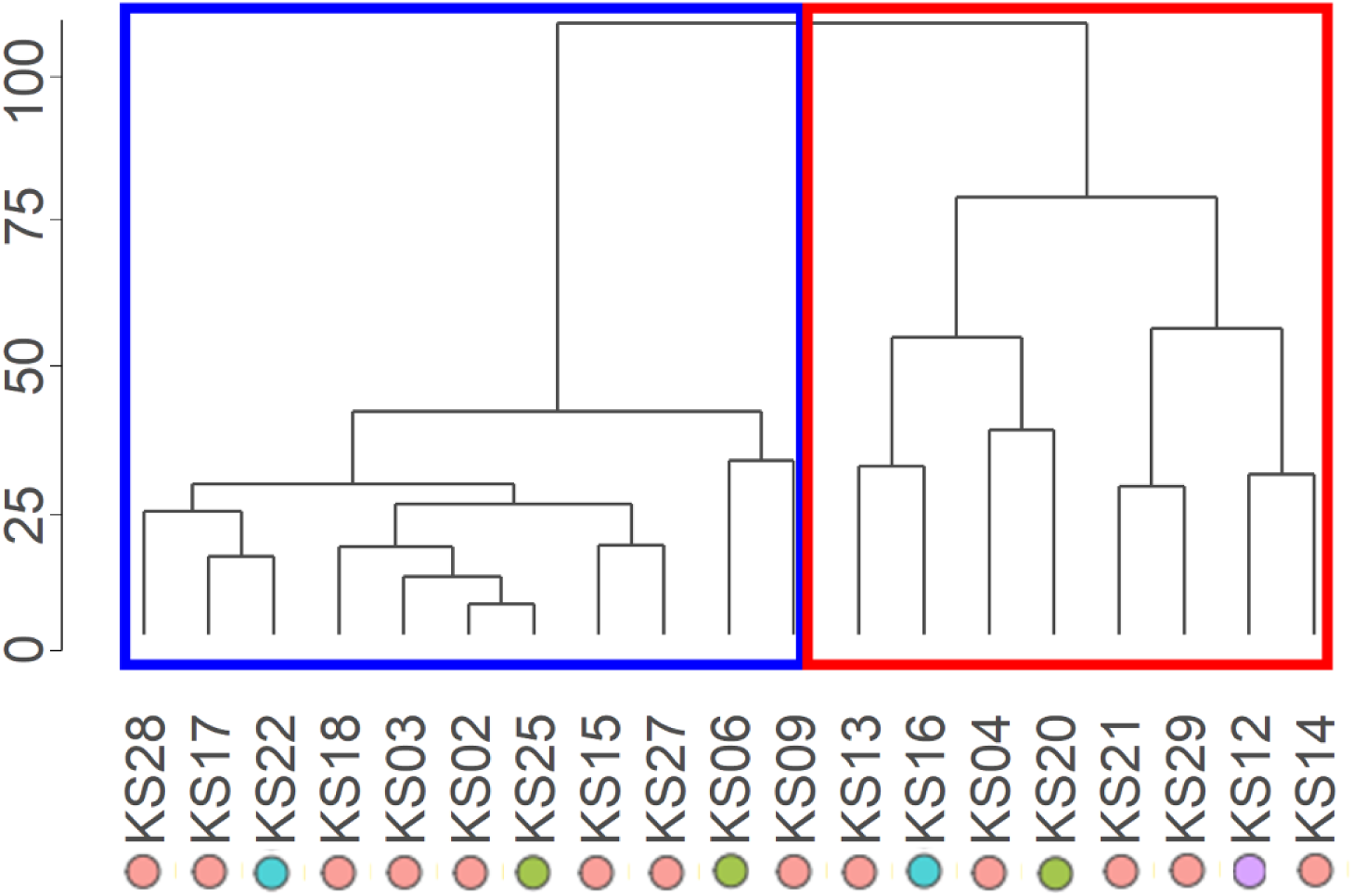
Native duckweed sites can be organised into derived from high light (dHL) or low light (dLL) using measured environmental light variables. Environmental light variables (n=45) measured in either µmol m^2^ s^-1^ (PPFD, FR, R, G, B, UV) or nm (λp, λd) for 19 sites across three seasons for two years were used to group sites by relatedness using a dendrogram. The distance matrix was computed using Manhattan distances and complete method. The rectangles represent site groupings after K means clustering set at n=2, where blue border indicates dLL sites and red border indicates dHL sites. Colored circles represent species groupings 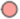 *L. minor*, 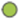 *L. minuta*, 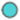 *L. turionifera*, 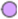 *S. polyrhiza*.

**Figure 4:**
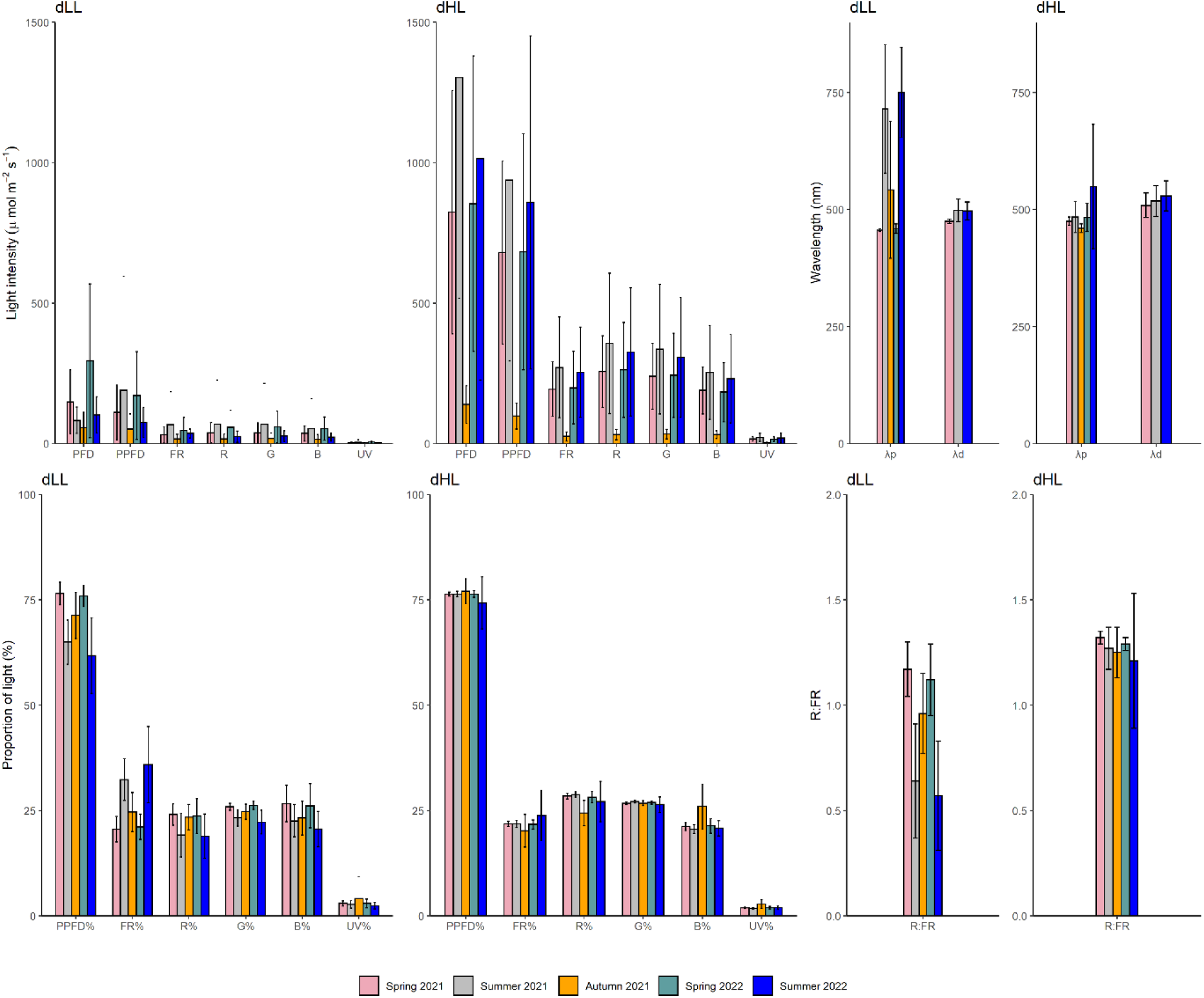
dHL and dLL sites have stark contrasts in photosynthetically relevant light environments. Light measured between different seasons and years from 19 duckweed sites grouped as dHL or dLL showing averages with error bars as ± standard deviation within site groupings. Light intensity and spectral quality was measured in µmol m^2^ s^-1^, for total light PFD – photon flux density is made up of regions of light FR – Far-red, R – Red, G-Green, B-Blue, UV-Ultraviolet and the photosynthetic portion grouped as PFFD – photosynthetic photon flux density (B+G+R). Wavelengths of light measured in nm were quantified for sites as λp – peak average wavelength and λd – dominant average wavelength. Proportions of spectral quality at each timepoint was calculated as proportion of spectral region/total PFD light x 100 per site and grouped by dHL and dLL. R:FR was calculated as ratio of R to FR light from raw values from each site and grouped as dHL or dLL. Bars are colored by seasonal and yearly timepoint in chronical order with dLL on the left and dHL on the right of each panel. Raw and grouped summary data and T test results are presented in Table S2.

Species were determined (see methods and Fig S5 Top panel). In general, individual ecotypes appeared to cluster into expected species groupings that corresponded with those from independent phenotypic assessment; of the 24 new UK ecotypes, four species were allocated including *L. minor* (n =16), *L. minuta* (n=5), *L. turionifera* (n=2) and *S. polyrhiza* (n=1). Although thought to be from the same species, KS16 and KS22 clustered separately from *L.turionifera* 9434, however their turiation (seed-producing) capacity led us define them as *L. turionifera*. Both *L.turionifera* 9434 and *S. intermedia* 9394 from the SRA had few genome reads giving severely low coverage <2% upon alignment. UK *L. minuta* and *S. polyrhiza* sequences generated from this study mapped significantly better with 10% coverage, whilst *L. minor* existing sequences and UK ecotypes both mapped with 80% and UK *L. turionifera* ecotypes mapped with 40% coverage, evidencing further species allocations by genetic similarity by mapping percentages. *L. minor* further clustered into two groupings of five and 11 individuals respectively, which could reflect a further degree of genetic variation amongst these ecotypes (Fig S5).

With regards to habitat, ecotypes of different *Lemna* species were found in both dHL and dLL environments, with a single *Spirodela* representative, *S. polyrhiza* from a dHL site (Fig 3, Fig 1D). There was high variation in seasonal growth patterns with *L. minor* accessions from a mixture of dHL and dLL sites which did not exceed 20% surface coverage across all timepoints of all years, remaining as sporadic colonies. As a contrast, *L. minor* ecotype KS03 from dLL had the highest average coverage overall, and maximum coverage in early Spring and Summer. The highest surface coverage achieved in different seasons/year time points were all from dLL sites, including KS06, KS02, KS25, KS18 showing superior maintenance of *L. minor* and *L. minuta* ecotypes in low light environments across extremes of seasonal light and temperature in the UK.

### Spatial and seasonal differences in light environments between sites

The key spectral differences between dHL and dLL sites following seasonal data collection are shown in Figure 4 (and Table S2). Light intensity and all spectral components were higher in dHL sites than dLL sites across all time points. Summer 2022 displayed the maximum recorded light intensity overall with greatest differences of PFD light levels: substantially lower in dLL (109 µmol m^2^ s^-1^) compared to dHL (1116 µmol m^2^ s^-1^). Peak wavelengths (λp) varied, with 750 nm (far-red) in dLL sites compared with 550 nm (green) light in dHL sites, especially in Summer. Dominant wavelengths between time points and dHL and dLL sites were similar around 500 nm. The proportions of spectral ratios across sites were also different with % FR and % UV higher in dLL sites and % R and % G higher in dHL sites. R:FR ratios were substantially higher in dHL sites consistent with natural canopy shading and presentyear-round, but with greatest differences in Summer. The % PPFD differed between sites, with dLL receiving less light in the photosynthetically active region in Summer and Autumn. Whilst differences in light intensity and spectral quality indicate marked difference in year-round habitats due to light, no significant differences were found between dHL and dLL sites for water and atmospheric temperature measured at each timepoint or for bioclimatic temperature and precipitation data from the Bioclimatic variables database (S3. Table).

### Habitat light environment determined growth rate in controlled conditions

Growth rate differences were derived from growth curves (Fig S5 Bottom panel) and used to find the difference between HL and LL response. The relationships between all growth rate parameters with each other are displayed in Figures 9, S8 and species and ecotype growth rate differences under light are shown in Figures S6 and S7.

Duckweeds showed higher growth in LL compared to HL through area gain and colony gain (ANOVA, *p* = <0.001) aswell as higher final fresh weight and freeze-dried weight (Figure 5). From all four growth parameters in the HL treatment, duckweeds from dLL sites had significantly higher growth rates than duckweeds from dHL but there was no difference between growth potential in low light (Fig 5). The ecotypes positively affected by HL treatment or with a less severely negative % change in growth rate, were from dLL sites and include *L. minuta* and *L. turionifera* species (Figure S7). The most productive ecotype across light conditions, was KS03, a dLL *L. minor* ecotype with maximum growth of 1 mm^2^ day^-1^ and the highest FW and FDW in HL and was still in the top five in LL (Figure S5).

**Figure 5.**
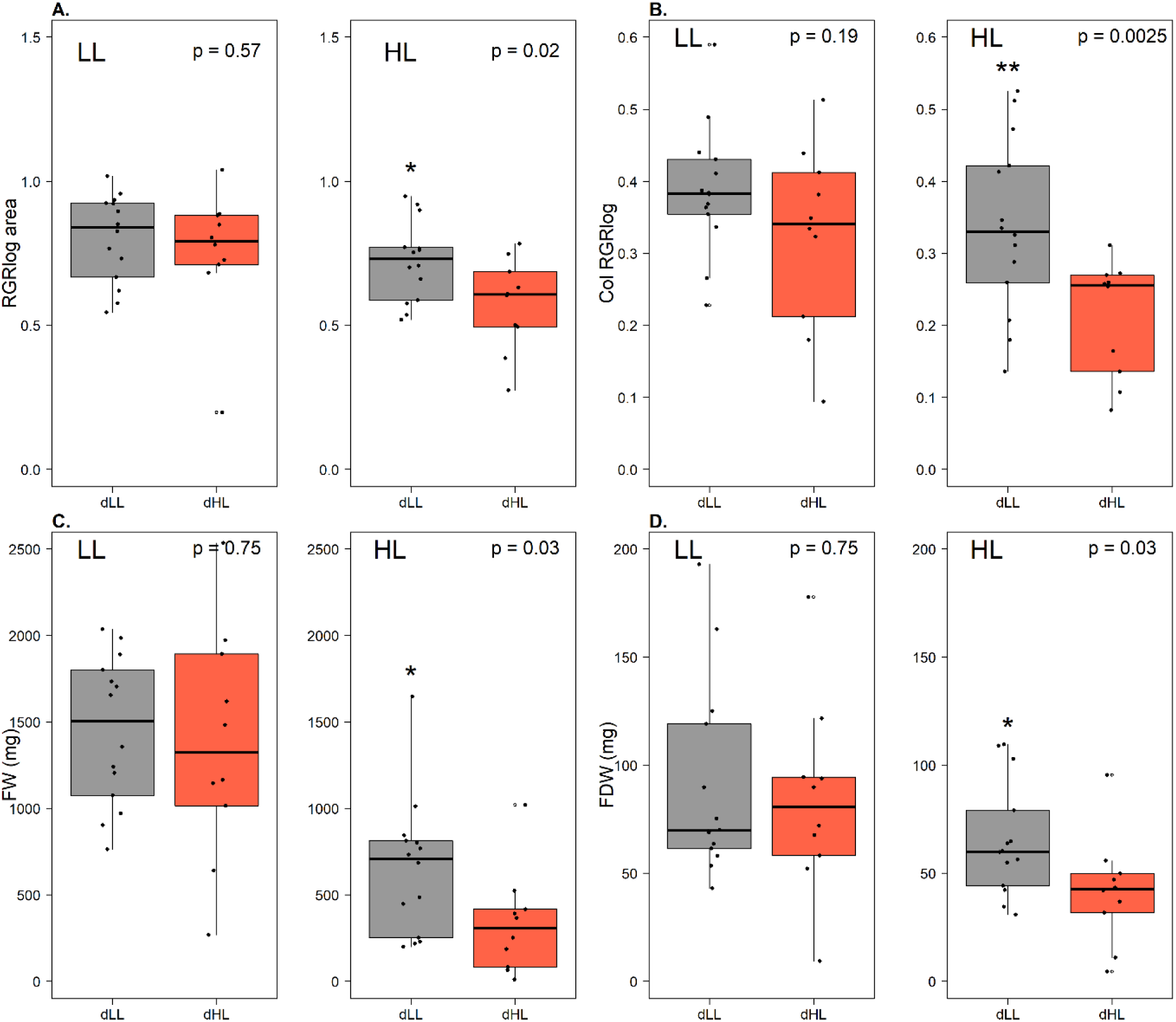
dLL ecotypes grow faster than dHL ecotypes in high light. Each pair of plots represent different growth rate parameters: A. RGRlog area (see Supplementary Fig 6A), B. Col RGRlog (see Supplementary Fig 6B), C. FW and D. FDW. Growth response in LL treatment on the left and HL on the right. Ecotypes are grouped as dLL (grey) or dHL (red). Midline on boxplots indicate median and 25% and 75% quartile boxes, outliers displayed as points. P values from Welch’s T-test are indicated above and significant differences by * <0.05 and ** <0.01.

### Photosynthetic efficiency in response to high light

Chlorophyll fluorescence imaging was used to see if the growth advantage by dLL plants was related to higher photosynthetic efficiency. Light response curves are presented for photosynthetic efficiency and NPQ for ecotypes grown in LL and HL treatments (Figure 6).

**Figure 6.**
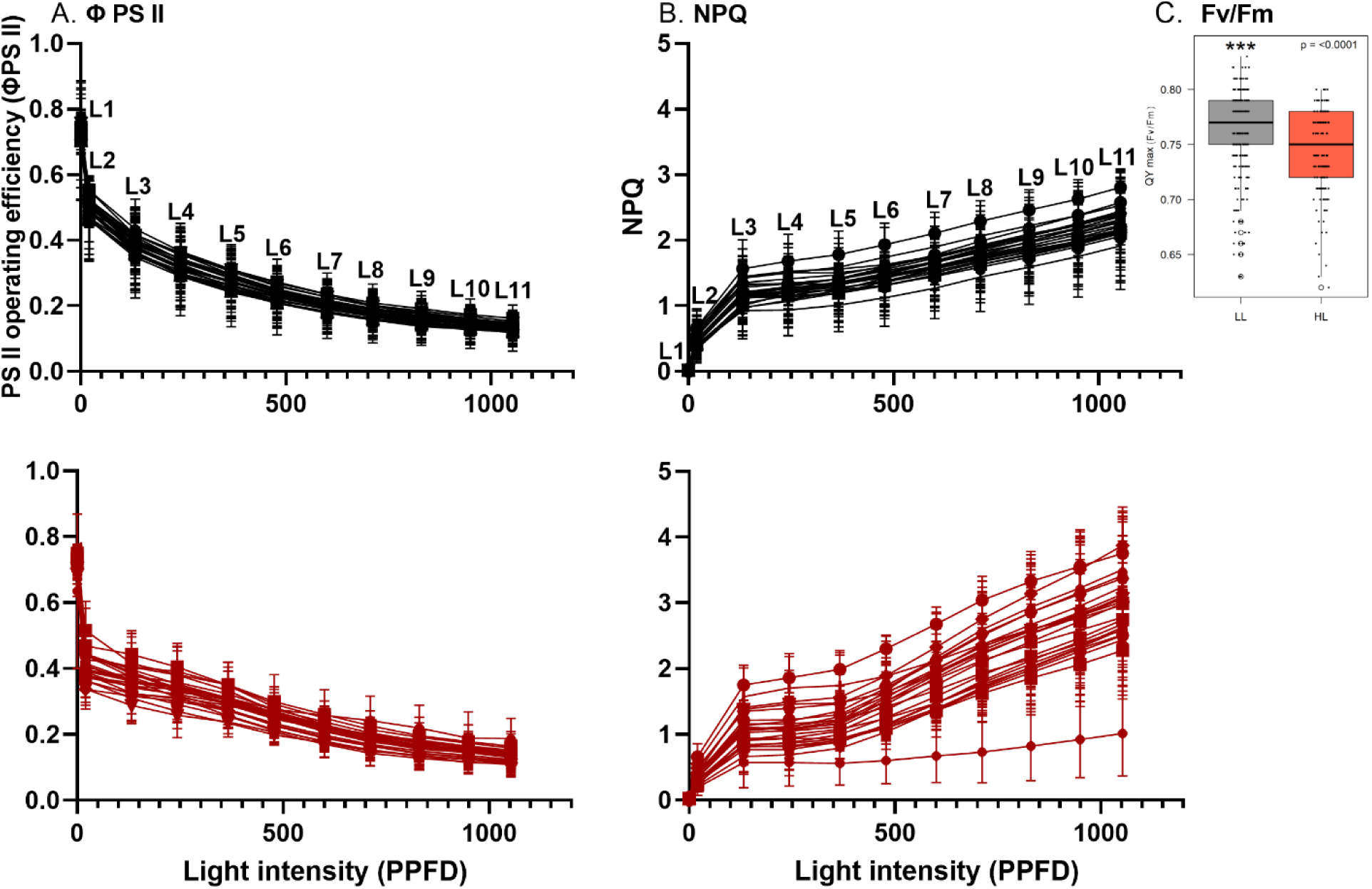
Photosynthetic efficiency (ϕ PSII) declines faster and higher maximum NPQ is induced in HL-grown plants in response to increasing light. *A.* ϕ PSII and *B.* NPQ measured at 11 light levels show average light response curves using Fluorcam and parameter calculations derived from (Murchie & Lawson, 2013). Dataset includes all ecotypes grown in LL (black) or HL (red) with n = 15 replicates for each treatment-ecotype combination grown from five separate growth rate experiments. Light levels allocated as L1-L11 correspond to PPFD: 0, 20, 130, 245, 365, 480, 600, 710, 830, 950, 1050 µmol m^-2^ s^-1^. The light level closest to growing treatment conditions for LL is L3 and HL is L5. Inset*: C.* Maximum photosynthetic quantum yield *F*_v_ */ F*_m_ **is higher in LL-treated plants than HL-treated plants**. Boxplot indicates variation in maximum *F*_v_ */ F*_m_ data calculated from chlorophyll fluorescence data across ecotypes measured in the dark after growth in LL (grey) or HL (red). *** P value = <0.0001 by Welch’s T test.

The maximum photosynthetic efficiency measured in the dark (*F*_v_/*F*_m_) and maximum NPQ at highest light level >1000 µmol m^2^ s^-1^ were highly affected by treatment, accession, species (ANOVA, *p* = <0.0001) and the interaction between factors (ANOVA, *p* = <0.001) for NPQ and (ANOVA, *p* = <0.01) for QYmax (Table 1, Figure 6), showing both genetic and environmental variation. Whilst all values were below 0.83, the lower values in HL grown plants indicates photoinhibition of plants under these conditions. As light levels increased, operational photosynthetic yield or **ϕ** PS II declined by 50% at LL (130 µmol m^2^ s^-1^) and a further 20-30% decrease by 350 µmol m^2^ s^-1^, depending on growth light level. NPQ increased with increasing light levels with a maximum at > 1000 µmol m^2^ s^-1^ (Figure 6). NPQ capacity was higher in the HL-grown plants and values of NPQ are affected by lower *F*_v_/*F*_m_ which is expected given the calculation based on Fm (Figure 6). Photosynthetic saturation (here defined by the lowest values in **ϕ** PS II) was less extreme in *L. turionifera* and concordant with higher % growth responses to HL (Figure S7), linking growth to photosynthetic efficiency maintenance in this case.

**Table 1.**
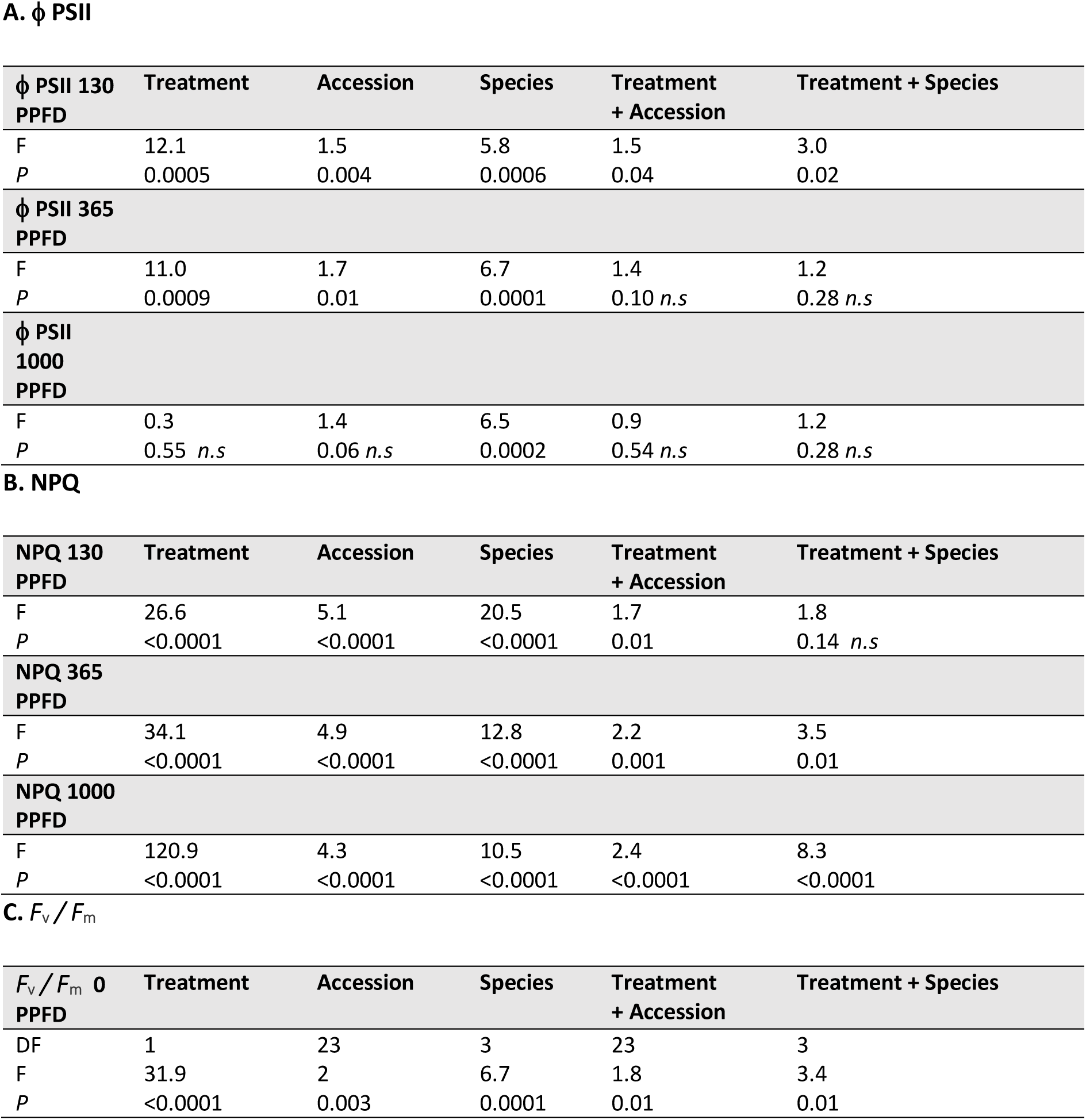
Photosynthetic efficiency (ϕ PS II), NPQ responses and quantum yield of photosynthesis (*F*_v_*/ F*_m_), vary between ecotypes, species and light treatments at corresponding low L3 (130 μmol m^-2^ s-^1^) and high light L5 (365 μmol m^-2^ s-^1^), however ϕ PSII at maximum light L11 (1000 μmol m^-2^ s-^1^) was only sensitive to species differences. At the maximum light level, only species had a significant effect on ϕPSII, as *L. turionifera* had higher ϕ PSII than *L. minor* and *L. minuta* grown in LL and HL at >1000 µmol m^2^ s^-1^ and at 130 µmol m^2^ s^-1^ and 365 µmol m^2^ s^-1^ light levels. Parameters derived from chlorophyll fluorescence. Table displaying Two-way ANOVA results using significance as <0.05. Non-significant results reported as *n.s*.

A. $\phi$ PSII
| $\phi$ PSII 130<br>PPFD | Treatment | Accession | Species | Treatment<br>+ Accession | Treatment + Species |
| --- | --- | --- | --- | --- | --- |
| F | 12.1 | 1.5 | 5.8 | 1.5 | 3.0 |
| P | 0.0005 | 0.004 | 0.0006 | 0.04 | 0.02 |
| $\phi$ PSII 365<br>PPFD | | | | | |
| F | 11.0 | 1.7 | 6.7 | 1.4 | 1.2 |
| P | 0.0009 | 0.01 | 0.0001 | 0.10 *n.s* | 0.28 *n.s* |
| $\phi$ PSII<br>1000<br>PPFD | | | | | |
| F | 0.3 | 1.4 | 6.5 | 0.9 | 1.2 |
| P | 0.55 *n.s* | 0.06 *n.s* | 0.0002 | 0.54 *n.s* | 0.28 *n.s* |

B. NPQ
| NPQ 130<br>PPFD | Treatment | Accession | Species | Treatment<br>+ Accession | Treatment + Species |
| --- | --- | --- | --- | --- | --- |
| F | 26.6 | 5.1 | 20.5 | 1.7 | 1.8 |
| P | <0.0001 | <0.0001 | <0.0001 | 0.01 | 0.14 *n.s* |
| NPQ 365<br>PPFD |  |  |  |  |  |
| F | 34.1 | 4.9 | 12.8 | 2.2 | 3.5 |
| P | <0.0001 | <0.0001 | <0.0001 | 0.001 | 0.01 |
| NPQ 1000<br>PPFD |  |  |  |  |  |
| F | 120.9 | 4.3 | 10.5 | 2.4 | 8.3 |
| P | <0.0001 | <0.0001 | <0.0001 | <0.0001 | <0.0001 |

**Table 1. Photosynthetic efficiency ( $\phi$ PS II), NPQ responses and quantum yield of photosynthesis ( $F_v/F_m$ ), vary between ecotypes, species and light treatments at corresponding low L3 (130 $\mu\text{mol m}^{-2} \text{s}^{-1}$ ) and high light L5 (365 $\mu\text{mol m}^{-2} \text{s}^{-1}$ ), however $\phi$ PSII at maximum light L11 (1000 $\mu\text{mol m}^{-2} \text{s}^{-1}$ ) was only sensitive to species differences.**
| $F_v/F_m$ 0<br>PPFD | Treatment | Accession | Species | Treatment<br>+ Accession | Treatment + Species |
| --- | --- | --- | --- | --- | --- |
| DF | 1 | 23 | 3 | 23 | 3 |
| F | 31.9 | 2 | 6.7 | 1.8 | 3.4 |
| P | <0.0001 | 0.003 | 0.0001 | 0.01 | 0.01 |

Higher *F*_v_/*F*_m_ in HL was associated with faster growth of ecotypes in both light conditions therefore photoinhibition may be the most important chlorophyll fluorescence component associated with high rates of growth in HL (Figure S8). Two/24 duckweed cultivars had increases in *F*_v_/*F*_m_ in HL relative to LL (Figure S7), this included *S. polyrhiza* KS12 and *L. turionifera* KS16. Whilst the higher *F*_v_/*F*_m_ values were in fast growing *L. minor* ecotypes and the ecotype with lowest *F*_v_/*F*_m_ in both HL and LL, and at various light levels was also the accession most drastically affected by HL in area gain, *L. minuta* LY01B (Fig S7A).

High NPQ was not directly associated with high growth in HL but instead linked to high growth in LL by area and fresh weight at 130 µmol m^2^ s^-1^ in LL and 130 and 365 µmol m^2^ s^-1^ in LL and HL grown plants. NPQ maxima were higher in *L. minor*, compared to other species, which also seemed better able to maintain *F*_v_/*F*_m_ indicating a reciprocal relationship between photoprotection and PSII efficiency.

To see whether ecotypes from light or shade environments had different photosynthetic and photoprotective responses in the light acclimation experiments, duckweeds were grouped by dHL or dLL. Unlike growth rate, no significant difference of habitat light level was recorded for *F*_v_/*F*_m_, NPQ or **ϕ** PSII at any measurement light level (L1-L11) by plants grown in HL or LL Thus habitat light environment did not directly relate to variation in photosynthetic parameters and photoacclimation.

Generally higher operational efficiency **ϕ** PSII or enhanced NPQ could not consistently be used to predict growth maintenance or improvement in HL and was not linked to previous differential light environments, so pigment composition was measured to see whether light-tolerant ecotypes maintained their photosynthetic pigmentation required for efficient light capture.

### Pigment rearrangements in high light adaptation are influenced by habitat light environment

The dLL ecotypes acclimated to HL by increasing carotenoid and maintaining chlorophyll to a greater extent than dHL ecotypes, as shown by higher Chl a, b total Chl (a+b) and carotenoids (Figures7, 8). There was no significant difference in Chl and Carotenoids between dHL and dLL ecotypes grown in the LL condition and there was no difference in overall Car:Chl ratios between dHL or dLL in either treatment (Figure 7). Light treatment affected pigment composition (ANOVA, *p* = <0.001, Table S9, Figure S10) as LL-grown plants had more Chl a and b, whilst HL-grown plants had more carotenoids and higher Car:Chl ratios indicating a photoprotective response overall (Figure 7). Although there were clear pigment rearrangements in responding to the light treatments in terms of total Chl and Car: Chl, duckweed ecotypes did not increase Chl a:b ratios to acclimate to HL and no differences was found between dHL and dLL ecotypes (Figure 7).

**Figure 7:**
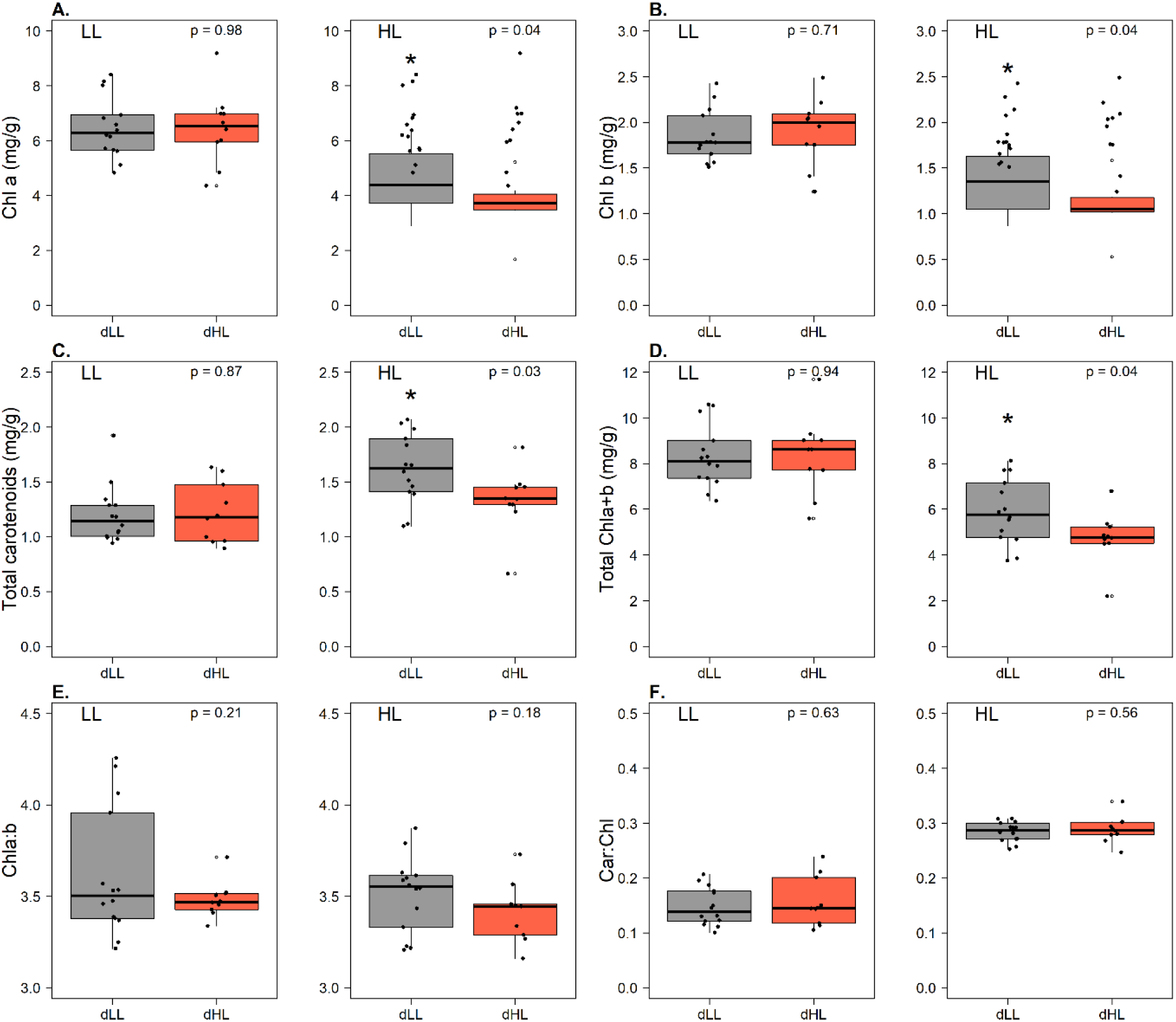
Chl a, b, total chlorophyll and carotenoid contents are higher in dLL duckweed ecotypes than dHL ecotypes when grown in HL but there are no differences between ratios of Chla:b and Car:Chl or any parameter in LL. Pairs of boxplots for pigment composition (mg/g) A. Chl a,B. Chl b, C. Total carotenoids, D. Chla+b and ratios E. Chl a:b and F. Car:Chl of ecotypes grown in HL and LL treatment. LL is on the left and HL on the right of each pair. Ecotypes are grouped and colored by dLL sites (grey) or dHL sites (red) on the X axis. Midline on boxplots indicate median and 25% and 75% quartile boxes. n= 20 for each ecotype-treatment combination from five separate growth rate experiments. All *P*-values by Welch’s T-test reported and all values <0.05 indicate significant difference.

Accessions from dLL sites lost less chlorophyll and had a greater increase in carotenoid content in HL (Figures 7, 8) than those from dHL sites (Figure 8B). The majority of dHL ecotypes had a greater loss in chlorophyll content when grown in high light and shifted towards the left of the plot (Figure 8A). *L. minor* KS13 was the only dHL ecotype to positively increase total chlorophyll in response to high light treatment.

**Figure 8:**
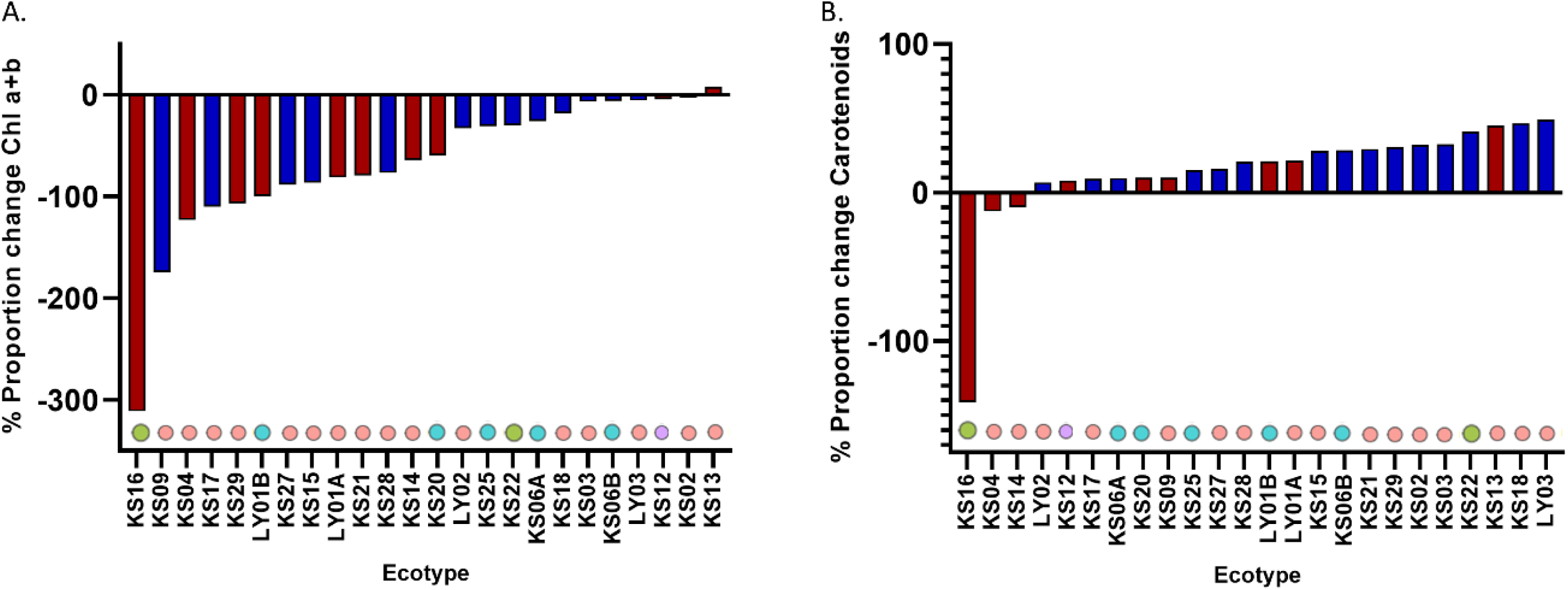
Total chlorophyll declines in response to high light whilst carotenoid content increases, with ecotypic variation in pigment contents. Difference in photosynthetic pigments in response to HL. *A.* % Proportional change in mean average Chl a+b difference between light treatments for each ecotype. *B.* % Proportional change in average total Carotenoid difference between light treatments for each ecotype. Barplots ordered on the X axis in descending order of ecotype response to HL treatment and colored according to original site light levels (red) dHL, (blue) dLL.

In a separate analysis, carotenoids were quantified using HPLC, (pooling leaf samples so that statistical analysis was not possible). Here, xanthophyll cycle (XC) carotenoids increased under HL (ANOVA, *p* = <0.0001), especially zeaxanthin (ANOVA, *p* = <0.0001), the intermediate antheraxanthin (ANOVA, *p* = <0.05) with a reduction in violaxanthin (ANOVA, *p* = <0.05). De-epoxidation state (DES) of XC increased in HL showing greater conversion to zeaxanthin in HL grown plants generally, with high levels of DES from 43% in LL to 67% in HL, .indicating relatively excessive light levels in both LL and HL in comparison to other higher plants under field conditions(e.g. Murchie *et al*., 1999)). No differences were found between dHL and dLL ecotypes for any carotenoid type (Figure S10). In this dataset we note Chl a/b was lower in LL grown plants overall, indicating acclimation of light harvesting complex antenna size may have occurred.

### Low light ecotypes adapted to high light by pigment rearrangements

The growth parameters measured in HL were highly correlated (Figure 9, Figure S8) and coincide with chlorophyll and carotenoid contents which contribute strongly to dataset variation (Figure 9). The Car:Chl ratio showed a strong negative correlation with RGR and pigment content in HL. In contrast to HL responses, pigment composition in LL apparently contributed little to growth rate in LL. Whereas the variables NPQ and *F*_v_/*F*_m_ acted in opposite directions. *F*_v_/*F*_m_ maintenance or increases in HL correlated positively with the HL growth response whilst NPQ responded positively with LL growth rate at various light levels (maximum > 1000 µmol m^2^ s^-1^, HL 365 µmol m^2^ s^-1^ and LL 130 µmol m^2^ s^-1^) (Figure 9, Supplementary Figure S8).

**Figure 9:**
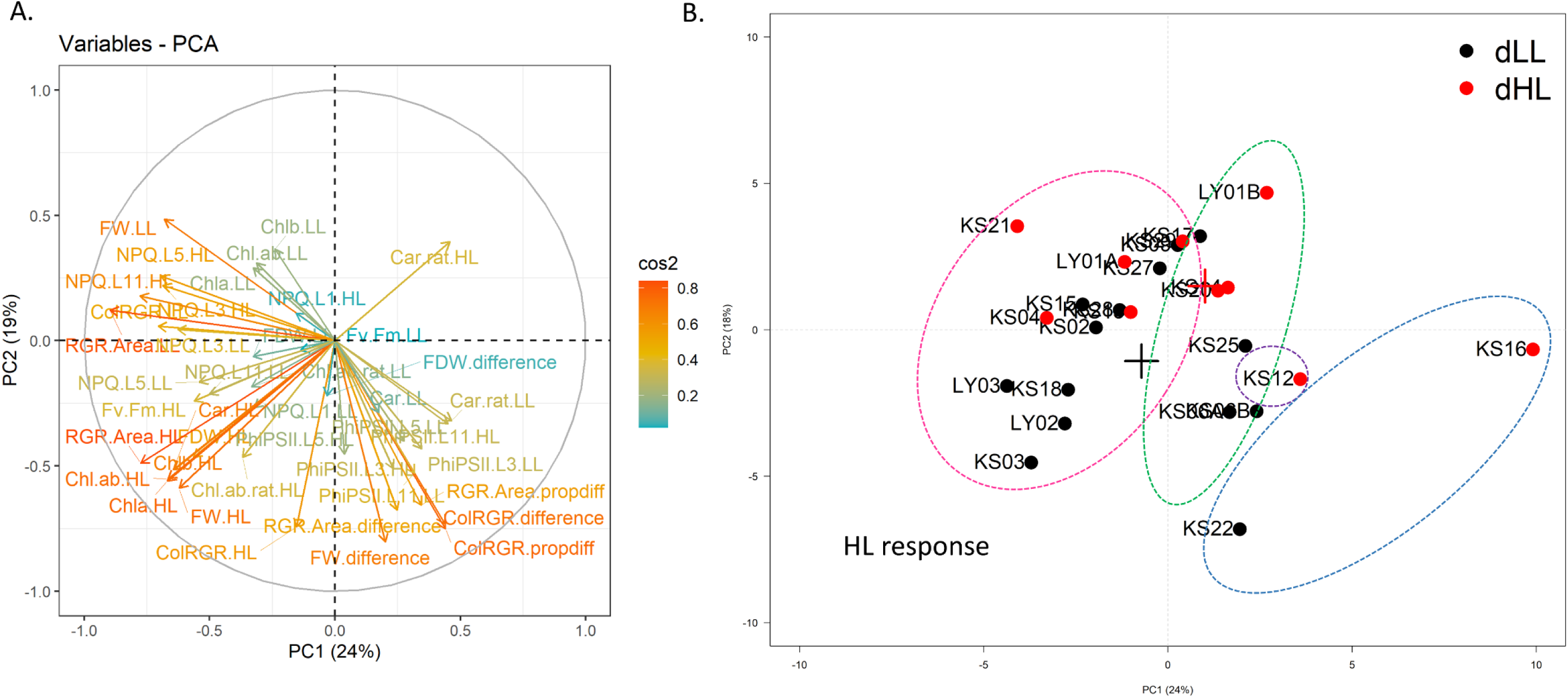
**A. Pigment composition correlates most strongly with growth parameters in HL, whilst LL growth is associated with NPQ responses at multiple light levels.** PCA showing association of physiological variables for 24 ecotypes grown in HL and LL conditions. Red cos2 values and long arrows indicate important contribution to dataset variation. Absolute and proportional % difference between HL LL response are included. **B. Duckweeds separate by HL-response and group into species when using light physiological responses with dLL diverging towards traits associated HL-performance and *L. minor* ecotypes variable in HL-responses.** PCA plot showing relationship between ecotypes in landscape of 42 physiological variables under high and low light treatment. Individual ecotype points are labelled and points are colored by dHL (red) and dLL (black) derived ecotypes. Centroids are marked with crosses for dHL and dLL ecotypes and indicate the average of values for all physiological variables in each grouping by original habitat. Ellipses represent clusters of species type 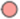 *L. minor*, 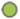 *L. minuta*, 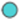 *L. turionifera*, 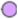 *S. polyrhiza*.

When ecotypes are plotted on the landscape of all physiological variables in light response, there is separation by environmental light groupings and species (Figure 9B). The fast growers in HL were all from dLL sites and comprise *L. minor* ecotypes KS03, LY02, LY03 and KS18, characterised by high pigment composition and high *F*_v_/*F*_m_ (Figure 9). The differences in mean centroids for dLL and dHL groupings, show separation by performance in HL primarily driven by variation in growth rate (Figure S5, S7) and pigment composition (Figures 7, 8). Species clusters in light response is also evidenced individually by significant differences for species on growth (ANOVA, *p* value = <0.0001, Table S6), photosynthetic parameters (ANOVA, *p* value = <0.0001 for NPQ and *F*_v_/*F*_m_, Table 1) and chlorophyll content (ANOVA, *p* value = <0.0001, Table S9).

## Discussion

Although a worldwide collective efforts of duckweeds has built an extensive collection available in stock centres (Sree & Appenroth, 2020) these are widely limited by unknown date of collection, specificity of collection locations and environmental data. Whilst shade tolerance and habitat is generally understood in higher plants, such information is less available for duckweeds and light adaptation remains less understood, with their aquatic habitat complicating simple categorisation. We generated a novel duckweed collection from diverse sites in North and South UK alongside detailed habitat data. We then explored variation in light acclimation in controlled environments. As well as ecophysiological understanding, such work is required for consideration of duckweeds and suitable growth environments for new food applications *e.g.* vertical farms and astrofarming (Hessel *et al*., 2022; Petersen *et al*., 2022).

### Colonisation in the field is linked to habitat light environment

Diversity in light quantity and quality in natural light environments is normally characterised in terms of PPFD depletion, scattering and green and FR enhancement caused by the presence of overhead vegetation (Lee, 1996; De Castro, 2000; Burgess *et al*., 2021). Stagnant aquatic habitats variably represent a slightly different case, with water edge and open water gaps in close proximity with colonization spreading between. Nonetheless, the differences between high and low light habitats represented largely expected features. Previously, low light sites characterized by tree shading were proposed to also provide temperature protection and contribute to higher nutrient injection into water from decaying biomass (Landolt, 1986; Landolt & Kandeler, 1987). Here, irradiance and spectral components were all significantly different between dHL and dLL sites year-round. Proportions of PPFD and R especially in Summer were more limited in dLL sites with higher FR. Shade environments are considered to limit light availability for photosynthesis and deplete photosynthetically active regions, however shade enriched components FR and green are still able to drive photosynthesis in well – acclimated plants (Smith *et al*., 2017; Zhen & Bugbee, 2020).

Supporting this, the highest plant coverage was maintained across dLL sites not dHL, showing ecotypes ability to acclimate to natural low light. Many dHL sites had sporadic growth and seemingly inhibited growth responses in many ecotypes. However it is important to note that no differences in temperature were noted between dHL and dLL sites so we conclude high light was an important driver of adaptation.

### Low light was more supportive for growth than high light in controlled conditions

Growth rates were generally higher in controlled low light (LL). Both low (< 100 µmol m^2^ s^-1^) and high light intensities have been cited as beneficial for biomass and growth rates in different duckweed species (Classen & Bergmann, 2000; Cheng *et al*., 2002) (Paolacci *et al*., 2018a; Stewart *et al*., 2020). Conclusions of species as a whole have been drawn from single clones of *L. minuta* and *L. minor* whilst a *L. gibba* clone was also grown in high light (Paolacci *et al*., 2018a; Stewart *et al*., 2020). However varied RGR measurement methods by area or biomass aswell as different starting densities, experimental duration and frequency of measurements challenge comparison of growth data between species across different studies. Optimal intensities for growth are still under question and here we show that they are also affected by collection origin. There was similarity of dLL and dHL accessions growth habits in LL treatment, which indicates both can grow well in low irradiance environments. In the HL treatment, growth was faster in dLL ecotypes. This observation is rather curious: perhaps dHL ecotypes had restricted growth rate due to local adaptation of high light. We hypothesize that a survival or stress tolerance strategy in dHL ecotypes was induced rather than promoting high growth. Similar established strategies can be seen for plants in nature (Pierce *et al*., 2017; Zhang *et al*., 2020). We suggest this is genetic in origin rather than epigenetic due to the multiple generations that took place between collection and experimentation, however it cannot be ruled out (Huber *et al*., 2021; Antro *et al*., 2022).

### Species variation of growth and photosynthesis

Variation among ecotype and species in growth rate has been previously shown with duckweeds grown in a common environment (Ziegler *et al*., 2015). Here we confirm species differences and now show a dependence on habitat origin. Previously, *L. minuta* were reported as faster growing than *L. minor* species (Ceschin *et al*., 2018) although here we refine this showing that *L. minor* ecotypes had the fastest growth rates in HL and LL but slower growing *L. minuta* and *L. turionifera* showed the greatest improvements in growth due to light.

The turionating species *L. turionifera (*KS16, KS22) and *S. polyrhiza* (KS12) were slow growers but enhanced/maintained growth in HL. Additionally, *L. turionifera* KS16 and *S. polyrhiza* KS12 were the only two cultivars which also had higher *F*_v_/*F*_m_ in HL (see below). As the only *Spirodela* ecotype, KS12 appeared to have additional mechanisms for photosynthetic acclimation to high light with a lower chlorophyll per dry weight, unchanged colony gain rate by treatment and no increase of (photoprotective) carotenoids induced by HL. In its wild habitat KS12 has high coverage in Summer and is visibly red (Figure 1D). The role of antioxidant mechanisms may be important drivers in species differentiation to light adaptation here, as anthocyanin and flavonoid genes are more expansive in *S. polyrhiza* and less prevalent in *Lemna* species (Landolt, 1986; Davies *et al*., 2022; Fang *et al*., 2023).

Generally, the photosynthetic response of *F*_v_/*F*_m_ and NPQ shown here is in line with studies of individual clones of *L. minor*, *L. minuta* and *L. gibba*, showing decreases in quantum yield and increases of NPQ in HL (Paolacci *et al*., 2018b; Stewart *et al*., 2020). In two UK duckweeds from dHL sites, atypical responses in HL occur with *F*_v_/*F*_m_ increasing in HL, *S. polyrhiza* and *L. turionifera*. Duckweed shows intra and inter-specific variation in photosynthetic parameters, as in other species including cereals (McAusland *et al*., 2019). While NPQ has a genetic basis underlying variation in rice (Kasajima *et al*., 2011), photoacclimation is also expected to have a genetic basis accounting for ecotypic and species variation seen here (Deblois *et al*., 2013; Lavaud & Lepetit, 2013; Tibiletti *et al*., 2016; Burgess *et al*., 2023).

### Maximum quantum yield (*F*_v_/*F*_m_) is closely linked to growth in HL

Phenotypic responses in plants are a product of both genome and environment (e.g. Ferguson *et al*., 2020). dHL ecotypes did not possess higher maximum NPQ than the dLL ecotypes, thus higher NPQ capacity was not required for HL. Relatively High NPQ in LL links it to high growth however the comparison with HL is made difficult by the lower *F*_v_/*F*_m_ which is intrinsically used for calculation of NPQ by comparing Fm with Fm’. Instead, *F*_v_/*F*_m_ correlated positively with higher HL growth and therefore is likely to contribute to the success of the dLL ecotypes in HL conditions. HL treatment caused a lower *F*_v_/*F*_m_ for 22/24 accessions, showing either photoinactivation / photoinhibition or perhaps zeaxanthin retention occurring in duckweeds. Thus it may be more important in this context to assess relative rates of damage to PSII and rate of repair as well as the ability to cope with irreversible damage. Recovery from photoinhibition is also promoted by other mechanisms such as antioxidant production which deserve further attention in duckweeds. Related to this flavonoid accumulation such as anthocyanins are induced by abiotic stresses in *S. polyrhiza* (Landolt, 1986; Böttner *et al*., 2021).

### Pigment responses

Chlorosis in duckweed can occur in response to light up to 1000 µmol m^2^ s^-1^ (Paolacci *et al*., 2018a), also associated with an increase in carotenoid levels (Stewart *et al*., 2020). Overall, plants from dHL lost more chlorophyll and carotenoid overall in HL than plants from dLL suggesting that chlorosis was also part of the adaption between sites. However we do not know if this was loss of chlorophyll per chloroplast, chloroplast number or cell structure.

Acclimation to HL is often accompanied by an increase in Car:Chl, as the xanthophyll cycle pool size increases and an increase in Chlorophyll a:b as antenna size reduces. Here, growth In HL did induce a higher Car:Chl across species and this also negatively correlated with growth rate, perhaps consistent with the importance of limiting PSII inactivation and the need for a higher *F*_v_/*F*_m_ for high growth rates in HL. Unchanged chla:b ratios have been shown for *Lemna* clones already (Paolacci *et al*., 2018a; Stewart *et al*., 2020) and other species such as barley in response to variable light (Murchie & Horton, 1997; Zivcak *et al*., 2014). Correlation of total chl a and b with HL growth rate indicates total chlorophyll retained is an important attribute for light adaptation. However chlorophyll a:b was inconsistent here, with the HPLC data showing a typical increase in HL. The HPLC data also showed high de-epoxidation rates in HL consistent with this being a highly light-saturating condition and further emphasizing the role of photoinhibition in determining growth. Overall it is likely a combination of acclimation and light induced stress in HL can be seen across our duckweed panel.

Reduced light intensity and extremely different peak wavelengths of light and altered proportions of spectral quality, including higher FR, reduced R and G, would be expected to affect light acclimation and chlorophyll content of ecotypes from low light habitats. Indeed, shade tolerant plants grown in controlled HL had higher chlorophyll and carotenoid contents per dry weight which correlate with HL growth, however measuring pigments per unit area would correlate better with photosynthetic capacity, as leaf area, water content and leaf thickness also tend to vary due to light acclimation (Lichtenthaler *et al*., 1981; Evans & Poorter, 2001). dHL ecotypes may be less sensitive to controlled high light, previously experiencing >1000 µmol m^2^ s^-1^ in their habitats and may acclimate in other ways, linked to a survival strategy rather than a dominant or competitor strategy, characterized by increased growth as already discussed. Related to this, we anticipate that the dHL conditions may have been hostile enough to induce a range of stress tolerance responses which we observe as *F*_v_/*F*_m_ reduction and pigment composition / reorganization.

## Conclusion

The habitat of duckweed ecotypes is important in selecting duckweeds for commercial purposes as environmental light contrasts were substantial and increased growth, chlorophyll retention and carotenoid gain was greater in dLL ecotypes in increased light conditions compared to dHL ecotypes. Further, we suggest that dHL ecotypes are not suitable for sustainable vertical farming systems. Subsequent breeding and genetic engineering options are limited in duckweed, highlighting the need to select cultivars by their environmental adaption and nutritional potential. The fastest growing ecotype with consistency was *L. minor* KS03, with high pigment contents, and may be a good ‘all-round’ option for commercial growth.

## Supplementary data

The following supplementary data are available.

**S1.** Data attachment. **Duckweed UK sites have four identified species and site coverage varies across season and year timepoints.**

**S2.** Data attachment. **Light variables used to characterise dHL and dLL groupings offer extreme light environments across seasons**.

**S3**. Data attachment. **Temperature and rainfall data across dHL and dLL sites indicate homogeneous conditions.**

**Fig S4. Negative relationship between duckweed surface coverage and Autumn UV light intensity.**

**Fig S5. Top. Clusters of Duckweed species differentiated by genome alignment and allele frequencies used to classify UK *Lemna* species using PCA. Bottom. Duckweeds show a period of lag followed by logarithmic growth rate patterns in green area and colony gain and RGR rates decline in HL-grown plants relative to LL-grown plants.**

**Table S6. Relative growth rate by area and colony gain vary between ecotypes and species in two common light treatment environments.**

**Fig S7. Top 2 panels: Growth by area and colony gain in duckweeds generally declines in response to high light treatment. Bottom 2 panels. The majority of duckweed ecotypes had increases in NPQ and decreases in *F*_v_/*F*_m_ in HL compared with LL treatment. dLL and dHL had no effect on photosynthetic parameters but species show differences.**

**Fig S8: Linear relationships between physiology of growth and photosynthetic responses in HL and LL treatments.**

**Table S9. Total Chlorophyll (a+b) varied between accession, species and light treatment but carotenoid levels only by light treatment and accession.**

**Fig S10. A. Carotenoid pigments separated by HPLC using chromatogram at 470 nm absorbance**. **B. Ratios of carotenoid pigment composition of HL and LL grown duckweeds vary when using HPLC.**

## Acknowledgements

Thanks to Levi Yant for supporting the UK duckweed collection, genotyping and providing clones from South UK. Thanks to Emma Jougla for help with pigment extraction. Thanks to Alex Burgess and Dylan Jones for advise on photography and image analysis for duckweed % coverage and growth measurements. Thanks also to Todd Michaels for providing the annotated *L. minor* 7720 sequence for species alignment. Thanks to Petra Ungerer and Sasha Ruban for performing carotenoid analysis with HPLC.

## Author contributions

KES performed the field collection, lab experiments and analysed the data. LC and BT aided with pigment extractions. LM wrote the code for Chlorophyll fluorescence experiments. MH wrote the code for genotype alignment to define duckweed species. EHM supported the project. KES and EHM conceived the study and wrote the manuscript with all authors contributing to the final manuscript.

## Data availability

The sequences for duckweed genomes in this panel are deposited as project XXX on SRA.

## Symbols and abbreviations

dHL: Derived from high light habitat.
dLL: Derived from low light habitat.
NPQ (*F*_m_–*F*_m_′)/*F*_m_′): Non-photochemical quenching, rate of heat loss from PS II.
LL: Low light.
HL: High light.
**ϕ** PSII (*F*_q_’/*F*_m_’): Operating efficiency of photosystem II electron transport in the light.
chla:b ratio: Ratio of Chlorophyll a to Chlorophyll b.
µmol m-2 s-1: Photons of light per metre per second.
PPFD: Photosynthetic photon flux density (400 – 700 nm).
PFD: Photon flux density (380 – 780 nm).
PFD-UV: Photon flux density UV light (between 380 – 400 nm).
PFD-B: Photon flux density Blue light (between 400 – 500 nm).
PFD-G: Photon flux density Green light (between 500 – 600 nm).
PFD-R: Photon flux density Red light (between 600 – 700 nm).
PFD-FR: Photon flux density Far-red light (between 700 – 780 nm).
λd: Dominant wavelength of light (color perceived).
λp: Peak wavelength of light intensity.
R:FR: Ratio of Red to Far red light.
GATK: Genome analysis toolkit.
SNP: Single nucleotide polymorphisms.
PCA: Principal component analysis.
RGR: Relative growth rate.
Col RGRlog (Col gain day-1): Relative growth rate during log phase by colony gain per day.
RGRlog area (mm-2 day-1): Relative growth rate during log phase by area gain per day.
FW: Fresh weight (wet biomass).
FDW: Freeze-dried weight of biomass.
L1-L11: Light levels 1:11 of chlorophyll fluorescence light program.
QYmax *(F*_v_ */ F*_m_): Maximum quantum yield of photosystem II photochemistry.
HPLC: High-performance liquid chromatography.
Chl a: Chlorophyll a.
Chl b: Chlorophyll b.
DEPs: De-epoxidation state of xanthophyll pool.
XC: Xanthophyll cycle pool
Car:Chl ratios: Ratio of carotenoids to chlorophyll content.

